# Central Nervous System Delivery and Biodistribution Analysis of an Antibody-Enzyme Fusion for the Treatment of Lafora Disease

**DOI:** 10.1101/671214

**Authors:** Grant L. Austin, Zoe R. Simmons, Jack E. Klier, Brad L. Hodges, Robert Shaffer, Tracy R. McKnight, James R. Pauly, Dustin Armstrong, Craig W. Vander Kooi, Matthew S. Gentry

## Abstract

Lafora disease is a fatal juvenile epilepsy, characterized by the malignant accumulation of aberrant glucan inclusions called Lafora Bodies (LBs). Cerebral delivery of protein-based therapeutics for the clearance of Lafora Bodies remain a unique challenge in the field. Recently, a humanized antigen-binding fragment (hFab) derived from a murine systemic lupus erythematosus DNA autoantibody (3E10) has been shown to mediate cell penetration and been proposed as a broadly applicable carrier to mediate cellular targeting and uptake. We report studies on cerebral delivery of VAL-0417, an antibody-enzyme fusion composed of the 3E10 hFab and human pancreatic α-amylase for the clearance of LBs in a mouse model of lafora disease. Herein, we report development of an enzyme-linked immunosorbant-based bioassay to detect VAL-0417 post treatment as a measure of delivery efficacy. We demonstrate the robust and sensitive detection of the fusion protein in multiple tissue types. Using our method, we measured biodistribution in different methods of delivery. We found intracerebroventricular administration provided the most robust delivery, while intrathecal administration only showed modest biodistribution. These data define critical steps in the translational pipeline of VAL-0417for the treatment of Lafora disease.

## Introduction

Lafora disease (LD, OMIM #254789) is an autosomal recessive fatal epilepsy that is also classified as a glycogen storage disease (GSD).^1–3^ There is currently no therapy, treatment, or cure for LD. Patients present with myoclonic, tonic-clonic, and focal occipital seizures caused by a build-up of abnormal, cytoplasmic glycogen-like aggregates called Lafora Bodies (LBs).^2–4^ LBs form in cells from nearly all tissues but are most insidious in neurons and astrocytes where they cause neurodegeneration.^5,6^ LD is the result of mutations in either *Epilepsy*, *Progressive Myoclonus type 2A* (*EPM2A*) or *EPM2B*, encoding the glycogen phosphatase laforin and the E3-ubiquitin ligase malin, respectively.^1,7–9^

LBs are comprised of glucose connected by α-1,4- and α-1,6-linkages and synthesized by the same enzymes as glycogen. However, the glucose chains are longer, less branched, and possess higher levels of covalent phosphate.^10,11^ These characteristics cause the molecules to become starch-like and aggregate into pathogenic LBs. Multiple labs have demonstrated that LBs drive neurodegeneration in both LD and non-LO models.^12–17^ Therefore, an immediate goal for LD treatment is to clear LBs and/or prevent their accumulation. Currently, multiple paths to inhibit LB formation and/or clear LBs once they are formed are being pursued.^18,19^ One therapeutic method utilizes antibody-enzyme fusions (AEFs), proteins comprised of an antibody domain fused to an enzyme by a linker peptide, that integrate the required functionalities to degrade LBs and penetrate cells.^20,21^

Cytosolic targeting for protein therapeutics is challenging, but recent antibody-based delivery platforms provide a unique opportunity to penetrate the cell membrane.^22–29^ The monoclonal anti-DNA autoantibody 3E10, produced in a murine model of systemic lupus erythematosus, has been shown to penetrate cells.^30^ Additionally, its antigen-binding fragment (Fab) and its variable domain (Fv) fragment can be fused to an enzyme and promote cytosolic delivery in multiple cell types.^25–29^ 3E10 fusions penetrate cells using a nearly ubiquitously expressed nucleotide salvage pathway mediated by the equilibrative nucleoside transporter 2 (ENT2, SLC29A2).^23,24,30,31^ Proteins with a large molecular weight, and those not able to traverse membranes on their own, enter cells expressing ENT2 when they are linked to 3E10 fusions.^25–27^ 3E10 alone is also an effective co-therapy against certain cancers when used with DNA damaging treatments, i.e. Doxorubicin or radiation^29^, and 3E10 has been shown to be safe in phase 1 clinical trials.^32^

The AEF VAL-0417 was recently created using a 3E10 hFab, capable of cell penetration and well-suited for clinical use, genetically fused with human pancreatic α-amylase, an enzyme that naturally degrades LBs.^33^ As a key tool to allow monitoring and iterative design of a dosage protocol, we developed an indirect sandwich enzyme-linked immunosorbent assay (ELISA) to specifically detect and monitor VAL-0417. The assay utilizes a capture antibody specific for the DNA binding region of VAL-0417 (anti-3E10 Fab) and allows for orthogonal detection of the fused amylase. The ELISA yields accurate quantitation of VAL-0417 delivered to tissues or circulating in the blood, even in tissues with high endogenous amylase activity. Furthermore, straightforward modifications to this ELISA will allow the development of ELISAs for other biologicals using the 3E10-based AEF delivery platform. Using this assay, we define the appropriate method of VAL-0417 central nervous system (CNS) delivery. Administration of VAL-0417 into the CNS via the spinal cord or subarachnoid space, i.e. intrathecal administration, resulted in only moderate detection in the brain and mostly in the caudal regions. Conversely, we find that administration directly into the brain ventricles by intracerebroventricular administration allows more efficient CNS delivery of VAL-0417.

## Experimental Section

### Western blot

Western blots were run using TGX stain-free pre-cast 10% gels (Bio-Rad 456-8034) with 1X running buffer [100 ml 10X buffer (30.30 g/L Tris Base, 144 g/L Glycine, 10 g/L SOS) in 900 ml water]. Transfers were performed using a semi-dry turbo transfer system onto PVDF membranes (Bio-Rad 170-4274). The membranes were blocked with 5% non-fat milk in TBS (60.55 g/L Tris Base, 87.66 g/L NaCl, pH 7.4). Primary antibodies [rabbit anti-pancreatic amylase (Abeam ab21156, lot GR283033-25) and anti-3E10 Fab (ab10A4, anti-id, Valerian Therapeutics)], were incubated with 5% non-fat milk in TBS for 1 hour at room temperature at 1/5000 and 1/500 dilutions, respectively. The anti-3E10 Fab antibody is a mouse monoclonal antibody against the DNA binding region of the 3E10 antibody, Fab, and Fv. It is similar to previously described anti-id antibodies against 3E10.^34^ Membranes were washed in TBS before being incubated with secondary antibody (Cell Signaling Technology goat anti-rabbit IgG, HRP-linked Antibody 7074, lot 27) at 1/7500 dilution. The membranes were washed, developed using ECL solutions (Bio-Rad 1705060), and imaged using a ChemiDoc MP imaging system (Bio-Rad 12003154).

### Enzyme-Linked Immunosorbant Assay

Wells of a Corning 9018 96-well plate were incubated with 100 μL of capture antibody (mouse anti-3E10 Fab, ab10A4, anti-id, Valerian Therapeutics) overnight (~16 hours) at a final concentration of 2 μg/ml in TBS at 4°C. All incubations were done in a humidified chamber. Wells were then rinsed (solution applied and then immediately removed) 3 times with 200 μL TBS followed by incubation for 1 hour with 200 μL of blocking solution (5% non-fat milk in TBS) at room temperature. Wells were then rinsed 3 times with 200 μL TBS. 100 μL of sample (tissue or recombinant protein sample diluted to 100 μL in TBS) was added and incubated for 1 hour at room temperature. Wells were rinsed 3 times with 200 μL of TBS and 100 μL of primary antibody (rabbit anti-pancreatic amylase, Abeam ab21156, lot GR283033-25) in 5% non-fat milk in 1xTBS was added and incubated for 1 hour at room temperature. The wells were washed (solution added and incubated for 5 minutes before being removed) 3 times with TBS. 100 μL of secondary antibody (Cell Signaling Technology goat anti-rabbit IgG, HRP-linked Antibody 7074, lot 27) in 5% non-fat milk in TBS was added and incubated for 1 hour at room temperature. Wells were washed 5 times with TBS. 100 μL of TMB substrate (Thermo Fisher N301, lot TA257953) was added and run according to the manufacturer’s instruction. The reaction was stopped by adding 100 μL of stopping solution (0.18 M H_2_SO_4_). Absorbance was read for at 450 nm using a BioTek Epoch2 microplate reader.

To detect the amount of anti-3E10 Fab bound to the plate, goat anti-mouse IgG, HRP-linked antibody (lnvitrogen 62-6520, lot SG253194) was used at a 1/5000 dilution and developed using TMB substrate.

### Amylase Activity Assay

Amylase activity assays were performed according to the manufacturer’s protocol (BioVision K711-100) Briefly, samples were homogenized in amylase assay buffer (BioVision, K711-100-1) and these homogenized samples were also used in the ELISA and BCA assays. Following homogenization, the cleared lysate was mixed with a reaction mixture containing ethylidene-pNP-G7 (the substrate; ethylidene capped heptaose linked to p-Nitrophenol) and a porcine glucosidase. The reaction is run for one hour with absorbance readings taken every minute at 405 nm. Activity was then determined using the slope generated from these readings. Exception is made for the assay performed with LD mouse lysate. For the LD mouse lysate, tissue lysate was diluted rather than cleared so that the amylases would not pellet with the LBs.

### BCA Assay

Protein concentrations of the samples were determined using the Pierce BCA Protein Assay Kit (Thermo Scientific 23227) according to the manufacturer’s protocol.

### Mouse treatment and tissue harvesting

All procedures were approved by the Institutional Animal Care and Use Committee (IACUC) as specified by the 1985 revision of the Animal Welfare Act.

From untreated wild type (WT) mice, blood was obtained via a sub-mandibular artery bleed to collect whole blood in EDTA coated tubes (BO Microtainer 365974) followed by cervical dislocation and cranial dissection to collect the brain, which was homogenized in BioVision amylase assay buffer. Blood was diluted to the appropriate volume in 0.25M EDTA in PBS to give a solution that was 80% whole blood.

WT mice were treated with 0.08 mg/day of VAL-0417 or PBS for 14 days via intracerebroventricular (ICV) and then sacrificed on day 15 or were treated with 0.08 mg/day of VAL-0417 or PBS for 14 days via intrathecal (IT) administration and then sacrificed on day 15. Briefly, ICV administration involved a craniotomy on the mouse and insertion of a cannula with osmotic pump (ALZET®; DURECT Corporation) into the target ventricle. The catheter was anchored and pump was deposited in a dorsal subcutaneous pocked and the mouse was closed. IT administration involved surgical exposure of the cisterna magna. A catheter was guided through the lumbar region caudally with a stylet. The catheter was connected to an osmotic pump (ALZET®; DURECT Corporation). The catheter was anchored and pump was deposited in a dorsal subcutaneous pocked and the mouse was closed. Brains were harvested and sectioned into 6 sections before being frozen in liquid nitrogen. Brain surgeries, implantation of catheters, sacrificing of treated animals, and brain sectioning was performed by Northern Biomedical Research (NBR). Samples from NBR were shipped on dry ice and homogenized in BioVision amylase assay buffer.

LD mice^14^ (*Epm2a*−/−) were administered VAL-0417, recombinant pancreatic amylase, or PBS via ICV administration. LD mice were treated with 0.08 mg/day of VAL-0417 or 0.04 mg/day recombinant pancreatic amylase for 28 days and then sacrificed on day 29. Brains were harvested and quickly sectioned into 6 sections before being frozen in liquid nitrogen. Brain surgeries, implantation of catheters, sacrificing of treated animals, and brain sectioning was performed by NBR. Samples from NBR were shipped on dry ice and homogenized in BioVision amylase assay buffer.

### Statistics

Statistical analyses were performed using GraphPad Prism version 7.03 using parameters from previous studies.^35,36^ Error is presented as Standard Error of the Mean (SEM), unless otherwise stated.

## Results

### Development and Optimization of a VAL-0417 ELISA

VAL-0417 is an AEF generated by fusing the genes for the heavy chain hFab fragment of antibody 3E10 and pancreatic α-amylase (Figure 1A). The fusion protein is co-expressed with the light chain of the hFab fragment and the two chains are coupled via disulfide bonds during post-translational processing. After *in vivo* administration, VAL-0417 cannot be distinguished from endogenous amylase using anti-amylase antibodies or amylase activity assays. Additionally, a mouse monoclonal anti-3E10 Fab antibody generated against the Fab DNA binding domain is not robust enough for accurate quantification of low hFab fusion concentrations in tissue lysates. To address these deficiencies, a sandwich ELISA combining an anti-Fab capture antibody that is specific for 3E10 and a polyclonal rabbit anti-amylase primary detection antibody that provides signal amplification was developed to provide both specificity and robust signal for VAL-0417 detection (Figure 1B).

**Figure 1:**
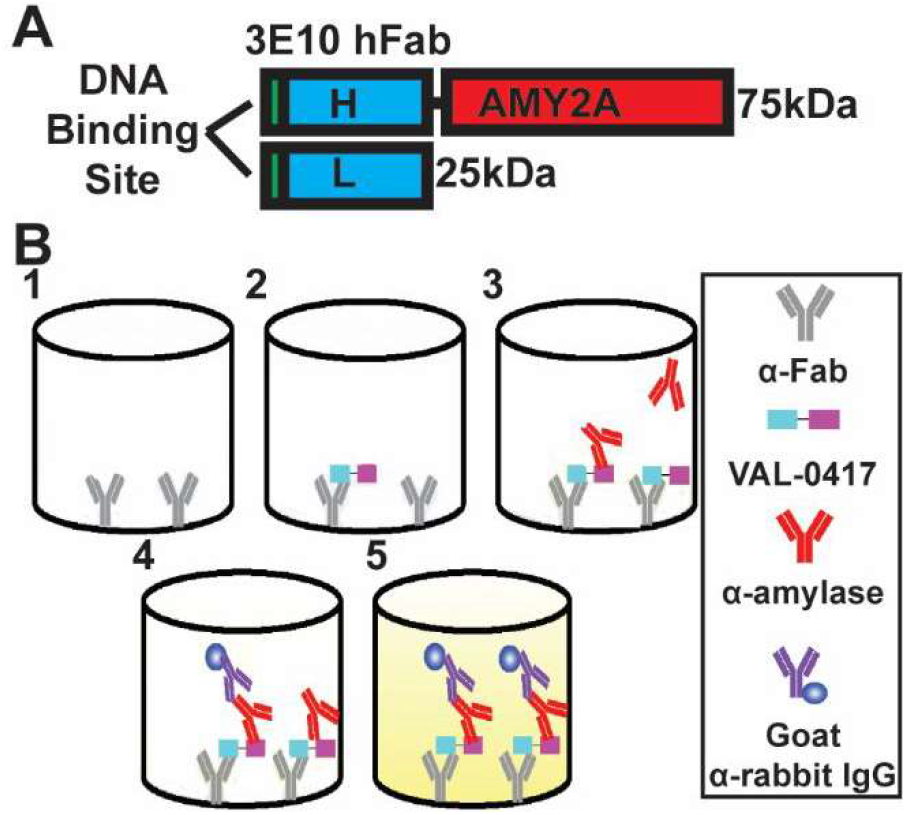
VAL-0417 and ELISA illustrations. **A.** Schematic of VAL-0417 demonstrating the hFab with its DNA binding site that is necessary for cell entry and the amylase protein that degrades LBs. H: heavy chain fragment of auto-antibody 3E10 Fab. L: light chain fragment of auto-antibody 3E10 Fab. AMY2A: human pancreatic α-amylase. **B.** Illustration of the ELISA method. 1: A plate is coated with the 10A4 anti-3E10 Fab capture antibody. 2: After blocking, wells are incubated with tissue lysate that may include VAL-0417. 3: VAL-0417 is detected with the Ab21156 rabbit anti-pancreatic amylase antibody. 4: The primary antibody bound is detected using a goat anti-rabbit HRP secondary antibody. 5: The amount of secondary antibody is detected by development with TMB substrate and H_2_SO_4_ to yield a yellow colorimetric product that is measured at 450nm.

It was first necessary to determine whether the proposed detection and capture antibodies would allow specific detection of VAL-0417. Recombinant VAL-0417, salivary amylase, and pancreatic amylase were resolved on a non-reducing SOS-PAGE, a Western transfer was performed, and the blot was probed with anti-pancreatic α-amylase (ab21156) or anti-3E10 Fab (ab10A4). The anti-pancreatic α-amylase antibody detected VAL-0417 and the two recombinant free amylases. Importantly, the anti-3E10 Fab antibody only detected VAL-0417 (Figure 2A). The anti-pancreatic α-amylase antibody detected two bands in the VAL-0417 lane, but the anti-3E10 Fab antibody only detected one band. This difference is because the anti-3E10 Fab antibody can only detect intact Fab that contains both the heavy and light chains whereas the anti-pancreatic α-amylase antibody detects both the protein and VAL-0417 without the Fab light chain. These data establish the utility of anti-3E10 Fab as a capture antibody and anti-amylase for detection.

**Figure 2:**
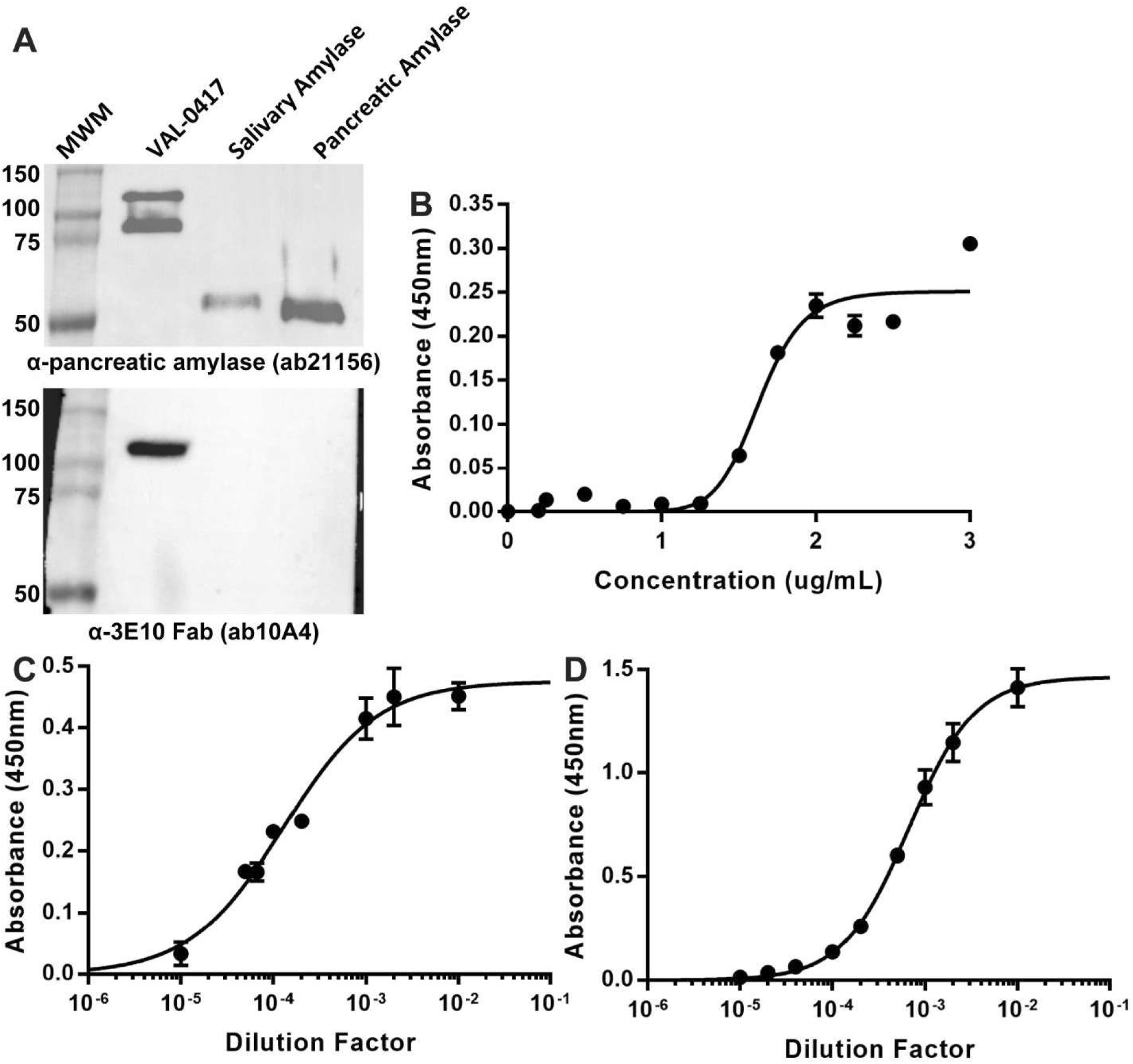
Optimization of antibodies for the VAL-0417 sandwich ELISA. **A:** Western blots using the rabbit anti-pancreatic amylase (top) and the anti-Fab (bottom). 1 μg of VAL-0417, salivary α-amylase, and pancreatic α-amylase were separated using a non-reducing SOS-PAGE, a Western transfer was performed, and the blot was probed with rabbit anti-pancreatic amylase (ab21156) or anti-3E10 Fab (ab10A4). Blots are representative from n=4. **B:** Determination of binding efficiency of the anti-Fab to a high binding plate. The capture antibody was plated onto a high binding 96-well plate and the amount of antibody adhered after overnight incubation was detected using a goat anti-mouse HRP antibody. The experiment uses n=3 replicates. **C**: Dilution curve of anti-pancreatic amylase antibody in the ELISA with constant concentrations of anti-Fab (2ng/μL), VAL-0417 (500 pg/μL), and goat anti-rabbit HRP secondary antibody (1/5000). The experiment uses n=3 replicates. **D.** Dilution curve of goat anti-rabbit HRP in the ELISA with constant concentrations of anti-Fab (2ng/μL), VAL-0417 (500 pg/μL), and anti-pancreatic amylase antibody (1/1000). The experiment uses n=3 replicates.

The concentrations of capture (anti-3E10 Fab) and detection (anti-pancreatic amylase) antibodies were then optimized for use in the proposed ELISA. Increasing amounts of anti-3E10 Fab were bound to wells of a high-binding 96-well plate from 0 ng/μL to 3 ng/μL to determine the concentration of maximally bound antibody. The anti-3E10 Fab antibody was maximally bound to the plate at a concentration of 2 ng/μL as determined by colorimetric analysis (Figure 2B). Optimization of the rabbit anti-pancreatic amylase and goat anti-rabbit IgG HRP detection antibodies was also performed. Titers of rabbit anti-pancreatic amylase and goat anti-rabbit IgG HRP detection antibodies were analyzed while maintaining constant concentrations of the capture antibody, antigen, and other detection antibody. The anti-pancreatic amylase and the goat anti-rabbit IgG HRP titers indicated a range of optimal combinations, i.e. any concentration within the linear range of the titer could be optimal (Figure 2C & 2D). Taking three points from the linear range of each titer gave nine possible antibody combinations that were tested using constant capture concentrations. These combinations were tested on high (H, 500 pg/μL), low (L, 10 pg/μL) and zero (0) concentrations of VAL-0417 and evaluated on the consistency of the replicates, the lack of background, the ability to detect the low concentration, and the signal range between the low and high concentrations (Figure S1). If results were equal, then the lowest antibody concentration was considered optimal. Cumulatively, these results demonstrated that the optimal antibody dilutions are 1/7500 for the anti-pancreatic amylase primary and 1/12500 for the goat anti-rabbit IgG HRP secondary.

The optimized antibody concentrations were then assayed using a series of VAL-0417 concentrations in triplicate ranging from 0.1 pg/μL to 10,000 pg/μL (Figure 3A). A best fit curve was generated to analyze the results using a four-parameter logistic^35^ (FPL) model (Equation 1). The FPL model was selected because it provides a better fit (percent recovery) to all of the data than the linear models, even in instances where the linear model lack-of-fit value (R^2^) is equal to or greater than 0.99.^35^

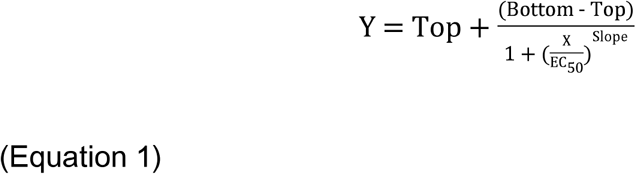

Y is the output value, in this case the absorbance at 450nm. Top is the upper asymptote and bottom is the lower asymptote.Xis the concentration of VAL-0417 in the well, EC_50_ is the concentration with 50% maximum response, and slope is the Hill coefficient.

**Figure 3:**
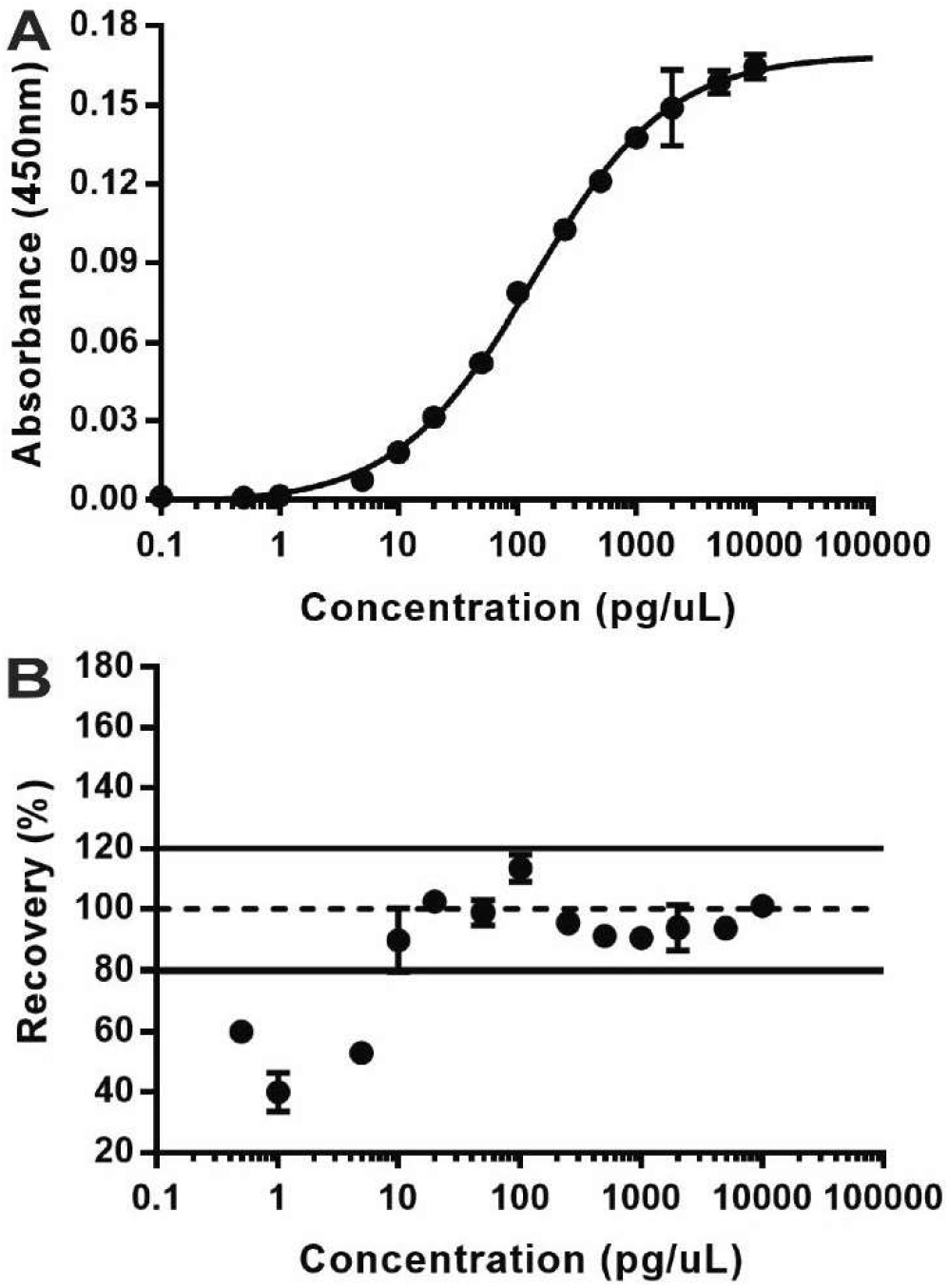
The working range of VAL-0417 in PBS at optimized antibody concentrations. **A**. An absorbance versus concentration curve was generated for multiple concentrations to determine the working range of the VAL-0417 ELISA. The minimum detection is 1 pg/μL and signal saturation is reached above 5000 pg/μL. The curve is fit with a four-parameter logistic (FPL) best fit curve. The experiment uses n= 3 replicates. **B.** Using the FPL model, percent recovery was determined by comparing concentration loaded versus the concentration as determined by the curve.

Evaluation of the percent recovery of each point with the concentration calculated by the FPL curve established that the lowest concentration measured within 20% of the FPL curve and that is significantly higher than background is 10 pg/μL or 10 ppm of VAL-0417 (Figure 3B). The upper limit is 5000 pg/μL, i.e. the point where the curve is saturated and absorbance no longer increases with increasing concentration. This model of standard curve with FPL fit is utilized to determine the concentration of VAL-0417 dose delivered.

### Tissue Matrix Effects on the VAL-0417 ELISA

Rapid and reliable analytical tests on whole blood are necessary for patient treatment and diagnosis, and they are important data in clinical trials. However, whole blood is an historically complex matrix for these assays.^37^ We sought to define the performance of the VAL-0417 ELISA in whole mouse blood to establish its reliability on samples in a complex matrix. Assays were performed to generate a standard curve in buffer versus 80% whole mouse blood (Figure 4). The slopes of the linear standard curves generated in the presence of blood were consistently decreased 3-fold versus standard curves in buffer (Figure 4A), yet they displayed a robust percent recovery throughout the majority of the standard curve when compared to the FPL curve (Figure 4B & 4C). The lower limit of detection increased to between 10 and 50 pg/μL. These data suggest that analysis of whole blood with this ELISA is viable when the standard curve is performed with appropriate referencing. Additionally, optimization of washing and incubation procedures within the ELISA protocol could help reduce these matrix effects. The tissue matrix effect was less pronounced when brain tissue was utilized as the matrix. When assays were performed with and without 10 μg protein from whole brain tissue homogenate or 100 μg protein from whole brain tissue homogenate, the ELISA standard curve difference between the slopes were 8.89% and 7.79%, respectively.

**Figure 4:**
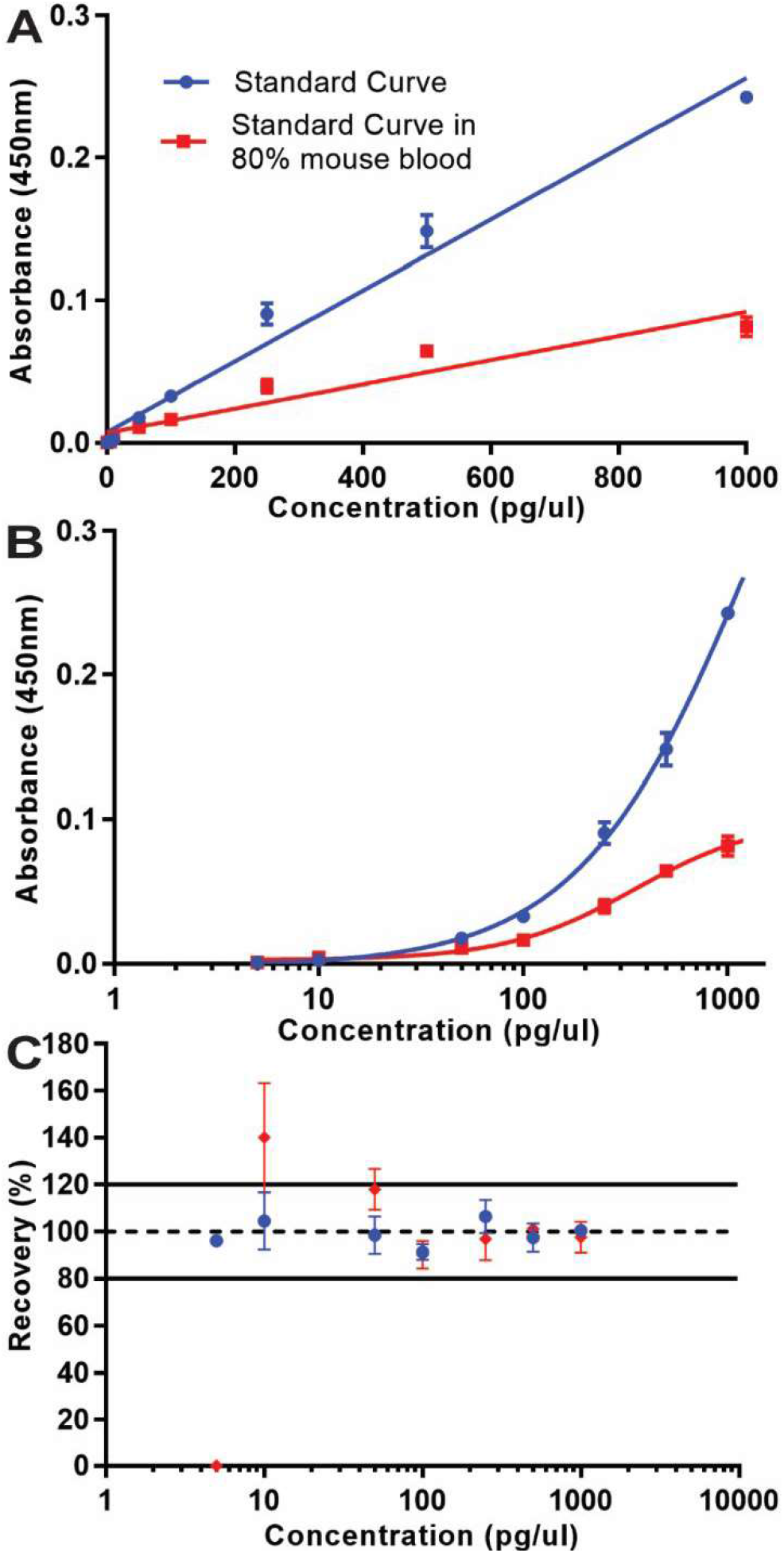
The effect of whole mouse blood on the VAL-0417 ELISA. **A.** A standard curve derived from varying concentrations of VAL-0417 in either PBS (Blue) or 80% whole mouse blood (Red) fit with a linear best fit curve. The experiment uses n= 3 replicates. **B.** The same standard curves as in **A** fit with an FPL best fit curve. **C.** Percent recovery of both curves against the FPL model best fit curve.

To test the specificity of the ELISA in the presence of tissue lysate, an *in vitro* screen was designed comparing equal mass concentrations of recombinant human pancreatic mylase and VAL-0417 with or without 50 μg protein from whole brain tissue homogenate. These parameters were chosen to test if the signal from biological samples is exclusively from VAL-0417 or if some signal is from endogenous amylases. The ELISA robustly detected VAL-0417 in the absence and presence of brain tissue homogenate and this signal was not changed by the presence of exogenously added pancreatic α-amylase. Conversely, there was no detectable signal from the sample with no antigen and no signal from the sample that only included pancreatic α-amylase (Fig. S2). Thus, the assay robustly differentiates between VAL-0417, pancreatic α-amylase, and endogenous amylases.

### Comparison of ICV and IT VAL-0417 Brain Delivery using the VAL-0417 ELISA

In order to define an optimal mode of therapy administration, initial pharmaco-delivery studies were performed on two modes of direct CNS drug delivery. Wild type (WT) mice were continuously infused with VAL-0417 (0.08 mg/day) or PBS for 14 days either via intracerebroventricular (ICV) administration or intrathecal (IT) administration. ICV administration is a method of direct infusion into the brain ventricles and IT administration utilizes the cerebrospinal fluid in the spinal canal to deliver treatment to the brain. ICV is more direct, but it is more invasive and requires brain surgery. One day after completion of VAL-0417 or PBS administration, the mice were euthanized, the brains were collected, and each brain sliced laterally into six sections labeled numerically from rostral to caudal (i.e. slice one was the most rostral portion of the cortex and slice six included the cerebellum, Figure 5 inset). In the ICV administered brains, VAL-0417 was detected in all six slices, although only significantly above background in slices 2, 3, and 4, with a maximal value of 1600 pg/μg protein (Figure 5). The administration site is within slice 3 and this is the same slice that yields the highest levels of VAL-0417. Conversely, IT administration did not yield high levels of VAL-0417 CNS distribution and the maximal ELISA detection was only 240 pg/μg protein (Figure 5). VAL-0417 was detected throughout the brain after IT administration, but the levels in all slices were dramatically less than with ICV administration after the same treatment length and no slice produced signal significantly higher than background. These results illustrate the utility of the VAL-0417 ELISA for pre-clinical analyses and the superior delivery via ICV administration compared to IT administration in mice.

**Figure 5:**
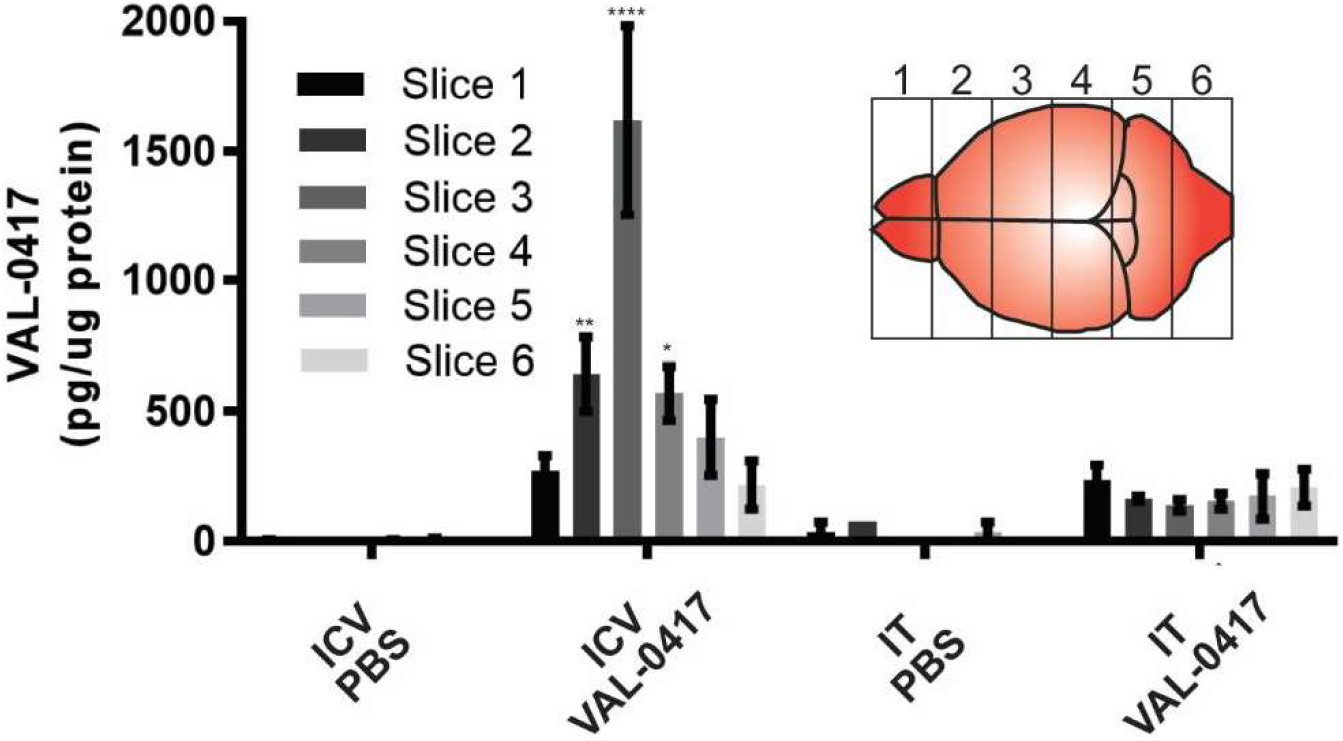
*In vivo* quantification of VAL-0417 in brain tissue after VAL-0417 treatment administered via intracerebroventricular (ICV) or intrathecal (IT) administration in WT mice. Distribution by slice of the concentration per μg of protein of VAL-0417 in the brain of WT mice after ICV or IT administration of PBS or VAL-0417 for 14 days. Mice were sacrificed one day following the completion of treatment (day 15) and brains harvested and sectioned laterally into six slices numbered from rostral to caudal (inset). These data include three mice in each group except the IT PBS group, which had two mice. Significance is measured by two-way ANOVA and is measured against the PBS measurement from the same slice and treatment duration. * is p≤0.05, ** is p≤0.01, and **** is p≤0.0001.

### CNS Delivery of VAL-0417 using ICV Administration in a LD Model

Finally, we sought to test the delivery efficacy of VAL-0417 and specificity of the ELISA in comparison with administration of amylase in vivo in an EPM2A−/− LD mouse model.^14^ LD is a fatal, neurodegenerative epilepsy and multiple groups have demonstrated that LBs in neurons and astrocytes drive disease progression.^12–17^ Therefore, VAL-0417 must be delivered to the central nervous system in order to be efficacious. LD mice were treated via ICV administration with PBS, or equal molar concentrations of VAL-0417 (0.08 mg/day) or recombinant pancreatic amylase (0.04 mg/day). After 28 days of continuous administration, the mice were euthanized, the brains were collected, and each brain sliced laterally into six sections labeled numerically from rostral to caudal. Robust levels of VAL-0417 were detected in slices 1-4 with the highest levels being those immediately adjacent to the administration site, i.e. slices two and three (Figure 6A), which is consistent with the ICV VAL-0417 distribution seen in the WT mice (Figure 5). Minimal to no signal was detected for the PBS treated or amylase treated animals in similar slices. These results indicate that VAL-0417 is disseminated throughout the brain during ICV administration and the ELISA robustly detects the AEF. While the ELISA is specific to VAL-0417, the amylase activity assay detects both VAL-0417 and any amylase. To confirm the ELISA results for VAL-0417 and determine distribution of the recombinant amylase, we performed amylase activity assays on samples from each brain slice. These results show similar trends as the ELISA and indicate that VAL-0417 is active in each slice and that the recombinant amylase without the hFab fused to it does not accumulate in the brain (Figure 6B). Therefore, the amylase assay confirmed the efficacy of the ELISA and the importance of the hFab fragment on VAL-0417 for delivery of amylase to the brain. While the amylase activity assay was important in this study to verify the delivery of recombinant amylase, the ELISA is a more optimal assay for the detection of VAL-0417. The ELISA provides less replicate variability, has a lower limit of detection, is more cost effective, and dramatically decreases background when compared to the amylase activity assay. Cumulatively, these results demonstrate that the VAL-0417 ELISA detects VAL-0417 after treatment, the ELISA differentiates between VAL-0417 and endogenous or exogenous amylases from treated samples, and ICV administration of VAL-0417 achieves robust biodistribution throughout the brain in the LD mouse model.

**Figure 6:**
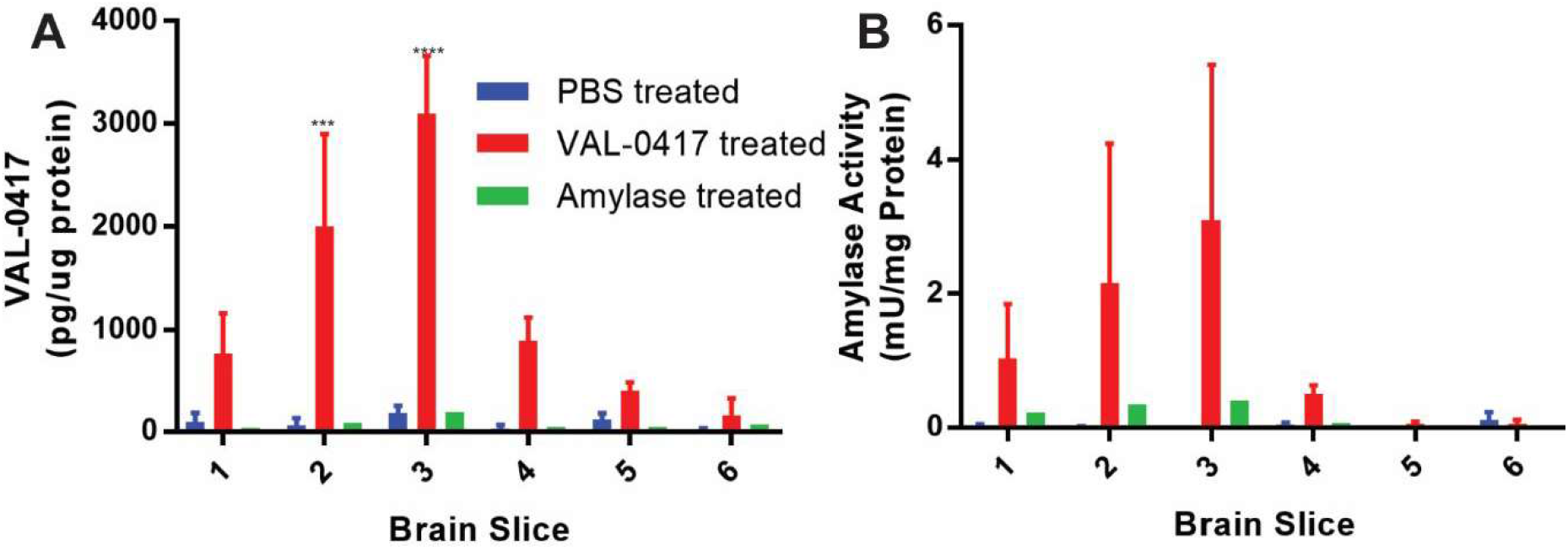
Determination VAL-0417 delivery in a LD mouse model. **A.** The concentration of VAL-0417 in brain tissue after treatment of LD mice by PBS, VAL-0417, or recombinant amylase as determined by the ELISA. Results are presented for each brain slice. **B.** Amylase activity in brain tissue after treatment of LD mice by PBS, VAL-0417, or recombinant amylase. This activity assay provides a confirmation of the ELISA results. Results are presented for each brain slice. These data include three mice each in the PBS and VAL-0417 groups and five mice in the Amylase group. Significance is measured by two-way ANOVA and is measured against the PBS measurement from the same slice. *** is p≤0.001, **** is p≤0.0001.

## Discussion

The development of quantitative detection assays for new therapeutics is an essential aspect of pre-clinical and clinical research. This study focuses on VAL-0417, a 3E10-based cell penetrating AEF that utilizes pancreatic amylase to degrade LBs and treat LD. We developed an ELISA for VAL-0417 and provide initial preclinical data demonstrating VAL-0417 delivery into WT and LD mice.

The sandwich ELISA, utilizing the anti-3E10 Fab and anti-pancreatic amylase antibodies, is a simple and robust method of detecting VAL-0417 concentration in samples from *in vivo* studies. The antibodies used in this ELISA function in combination to produce a robust signal that is specific for VAL-0417, which neither antibody could generate independently. The mouse monoclonal anti-3E10 Fab is highly specific for the DNA binding region of 3E10 and its fragments, but only produces a weak signal when used alone in an antigen-down ELISA format. The polyclonal rabbit anti-pancreatic amylase yields strong detection of amylases and also detects endogenous amylases. Furthermore, due to the interest in using VAL-0417 and related fusions for systemic delivery, these data demonstrate the effect of tissue matrices on VAL-0417 detection. Tissue matrix effects were found to result in systematic effects which can be accounted for with proper assay controls.

This ELISA is a substantial improvement to other methods that could be used for VAL-0417 quantification such as amylase activity assays, western blots, or mass spectrometric approaches. While amylase activity in brain is low, background signal would be a major issue in tissues of the gastrointestinal tract where endogenous amylase activity is high. This issue could be relevant in LD treatment because LBs are found throughout the body including high levels in the liver.^38^ It is currently unclear what the physiological effects of these systemic LBs are because patients die from their severe neurological symptoms before system effects are observed. These effects may become pronounced and could require intervention after patients are treated for their brain LBs via direct CNS therapy delivery. The ELISA is also superior because whole tissue lysate can be tested, whereas the amylase assay utilizes either dramatically diluted whole lysate or centrifugally cleared lysate. If the sample must be dramatically diluted, then the lower limit of detection will be increased. Clearing the lysate is also an issue in LD models because the VAL-0417 and all amylases would bind and pellet with LBs.

Two methods of VAL-0417 delivery to the brain were evaluated. While less invasive, the IT mode of administration did not yield as robust of a biodistribution of VAL-0417 throughout the brain in the mouse models. Conversely, ICV administration produced a robust signal after 14 days of administration. While ICV drug administration is invasive, it is already a relatively common procedure in the clinic.^39,40^ ICV administration of cerliponase alfa for children with CLN2 (neuronal ceroid lipofuscinosis type 2) disease is well tolerated and is becoming the standard of care.^41,42^ Currently, ICV administration is the optimal option for treating devastating neurological diseases like LD. However, novel methods are being developed to deliver biologics across the blood brain barrier (BBB) via noninvasive methods. One mechanism utilizes a modified Fe antibody fragment to interact with BBB transferrin receptors to facilitate transport via receptor mediated transcytosis.^43^ These emerging technologies provide exciting possibilities for future treatment of neurological diseases with AEFs such as VAL-0417.

LBs are most prevalent in the cerebral cortex, substantia nigra, and thalamus, which would correspond to slices 3, 4, and 5 in this study, and they are also present in the cerebellum, which would be slice 6 (Figure 5).^38^ The ICV administration site in this study was within a ventricle in slice 3 and the ICV administration achieved a maximum delivery load in this slice. VAL-0417 diffused in roughly Gaussian diffusion through the tissue from the injection site throughout the entire brain. Thus, ICV administration delivered VAL-0417 to the areas where it would be most needed.

More time dependent studies are necessary to elucidate the optimal delivery regimen of VAL-0417. The presented studies included dosing durations of two and four weeks, Moving forward, shorter treatment durations as well as intermittent treatment delivery should be tested. As with all therapeutic interventions, it will be necessary to determine the minimum effective dose and treatment duration for ICV administration before transitioning this therapy to the clinic. Furthermore, while the IT delivery was not optimal in this study, if the dose needed for therapeutic benefit is found to be lower, then IT administration may be the optimal choice due to the less invasive nature. While IT administration was not optimal in the mouse model, further testing in larger vertebrate models, including rats, should be explored. Additionally, both administration methods should also be evaluated in non-human primates to determine the optimal delivery method for the clinic. The ELISA method established here will be essential for all future studies evaluating the delivery and pharmacokinetics of VAL-0417.

The results presented herein are encouraging for the use of VAL-0417 as a LD therapy, which currently has no therapy or treatment. These data begin to establish the necessary delivery for VAL-0417 and will guide clinical trials designed to establish the efficacy of VAL-0417 to breakdown LBs in tissue and alleviate the devastating symptoms of LD.

## Supporting Information

Supplemental Figure 1: Screen to determine the optimal antibody concentrations for the VAL-0417 ELISA. (PDF)

Supplemental Figure 2: Specificity of the VAL-0417 ELISA with and without the presence of tissue matrix. (PDF)

## Author Information

Department of Molecular and Cellular Biochemistry, University of Kentucky College of Medicine, Lexington, KY 40536, USA. phone: 859.323.8482

The manuscript was written through contributions of all authors. All authors have given approval to the final version of the manuscript.

## Funding Sources

This work was supported by a sponsored project from Valerian Therapeutics to M.S.G. as well as NIH grants R01 NS070899, P01 NS097197, and an Epilepsy Foundation New Therapy Commercialization Grant all to M.S.G.

## Acknowledgements

The authors would like to thank Dr. Ramon Sun regarding initial conversations for the conception and development of the VAL-0417 ELISA. We would also like to thank Lyndsay Young, Kit Donohue, Bobby Murphy, and the rest of the Gentry lab for helpful discussions and assistance during manuscript preparation.

