## Supplemental Figures for "Central Nervous System Delivery and Biodistribution Analysis of an Antibody-Enzyme Fusion for the Treatment of Lafora Disease"

| | Secondary | 1 | 2 | 3 | 4 | 5 | 6 | 7 | 8 | 9 | 10 | 11 | 12 |
|---|---|---|---|---|---|---|---|---|---|---|---|---|---|
| Primary | | Blank | | 1/25000 | | | 1/12500 | | | 1/6250 | | 0,1/12500 | Blank |
| A | Blank | | | | | | | | | | | | |
| B | 1/30000 | | H | L | 0 | H | L | 0 | H | L | 0 | H | |
| C | | | H | L | 0 | H | L | 0 | H | L | 0 | L | |
| D | 1/7500 | | H | L | 0 | H | L | 0 | H | L | 0 | 0 | |
| E | | | H | L | 0 | H | L | 0 | H | L | 0 | H | |
| F | 1/1875 | | H | L | 0 | H | L | 0 | H | L | 0 | L | |
| G | | | H | L | 0 | H | L | 0 | H | L | 0 | 0 | |
| H | Blank | | | | | | | | | | | | |
| | | | | | | | | | | | | 1/7500,0 | |

| | Secondary | 1 | 2 | 3 | 4 | 5 | 6 | 7 | 8 | 9 | 10 | 11 | 12 |
|---|---|---|---|---|---|---|---|---|---|---|---|---|---|
| Primary | | Blank | | 1/25000 | | | 1/12500 | | | 1/6250 | | 0,1/12500 | Blank |
| A | Blank | | | | | | | | | | | | |
| B | 1/30000 | | 0.05 | 0.046 | 0.045 | 0.058 | 0.045 | 0.045 | 0.133 | 0.046 | 0.046 | 0.045 | |
| C | | | 0.058 | 0.045 | 0.045 | 0.056 | 0.045 | 0.045 | 0.123 | 0.045 | 0.045 | 0.045 | |
| D | 1/7500 | | 0.082 | 0.045 | 0.045 | 0.09 | 0.048 | 0.046 | 0.245 | 0.046 | 0.045 | 0.044 | |
| E | | | 0.068 | 0.044 | 0.044 | 0.085 | 0.048 | 0.045 | 0.147 | 0.046 | 0.045 | 0.044 | |
| F | 1/1875 | | 0.141 | 0.046 | 0.046 | 0.156 | 0.048 | 0.046 | 0.36 | 0.05 | 0.047 | 0.044 | |
| G | | | 0.126 | 0.045 | 0.046 | 0.242 | 0.048 | 0.046 | 0.502 | 0.053 | 0.05 | 0.045 | |
| H | Blank | | | | | | | | | | | | |
| | | | | | | | | | | | | 1/7500,0 | |

**Supplemental Figure 1:** Screen to determine the optimal antibody concentrations for the VAL-0417 ELISA. Determination of optimal detection antibody concentrations calculated from the curves in Figure 2C and 2D. The screen was run in duplicate using high (H, 500 pg/µL), low (L, 10 pg/µL), and zero (0, 0pg/µL) concentrations of VAL-0417. There were nine combinations with a negative control for each antibody for a total of 11 antibody combinations. Anti-3E10 Fab capture antibody was at a constant concentration (2ng/µL). Primary is the rabbit anti-pancreatic amylase and secondary is the goat anti-rabbit IgG HRP.

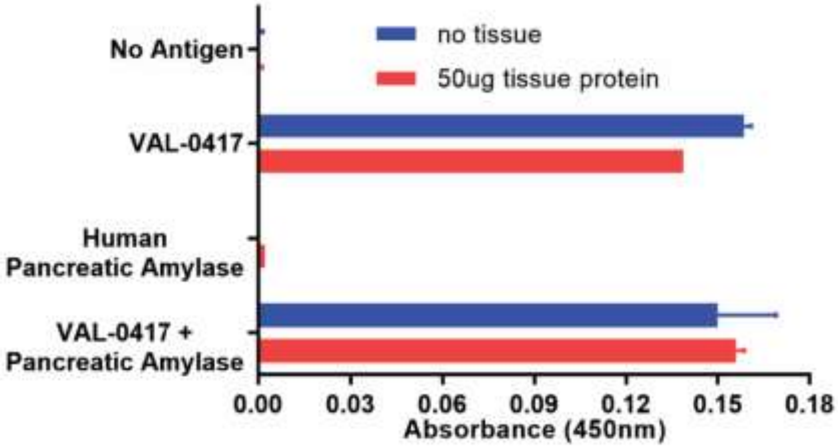


**Supplemental Figure 2:** Specificity of the VAL-0417 ELISA with and without the presence of tissue matrix. The ELISA was performed in triplicate using samples treated with VAL-0417, recombinant human pancreatic amylase, or both proteins at 250 pg/μL in the presence or absence of 50 μg homogenized mouse brain tissue protein.
